# DNA methylation associates with sex-specific effects of experimentally increased yolk testosterone in wild nestlings

**DOI:** 10.1101/2024.08.12.607609

**Authors:** Bernice Sepers, Suvi Ruuskanen, Tjomme van Mastrigt, A. Christa Mateman, Kees van Oers

## Abstract

Maternal hormones can profoundly impact offspring physiology and behaviour in sex-dependent ways. Yet little is known on the molecular mechanisms linking these maternal effects to offspring phenotypes. DNA methylation, an epigenetic mechanism, is suggested to facilitate maternal androgens’ effects. To assess whether phenotypic changes induced by maternal androgens associate with DNA methylation changes, we experimentally manipulated yolk testosterone levels in wild great tit eggs (*Parus major*) and quantified phenotypic and DNA methylation changes in the hatched offspring. Increased yolk testosterone levels decreased the begging probability, emphasised sex-differences in fledging mass and affected methylation at 763 CpG sites, but always in a sex-specific way. These sites associated with genes involved in growth, oxidative stress and reproduction, suggesting sex-specific trade-offs to balance the costs and benefits of exposure to high yolk testosterone levels. Future studies should assess if these effects extend beyond the nestling stage and impact fitness.

## Introduction

Environmental conditions experienced by mothers can have profound impacts on the phenotypes of their offspring (Mousseau and Dingle, 1991; Bernardo, 1996; Kofman, 2002; Groothuis et al., 2005; Maestripieri and Mateo, 2009). These so-called environmental maternal effects are aspects of the maternal phenotype that cause changes in offspring phenotype and have been documented across a wide array of taxa (e.g. Räsänen and Kruuk, 2007). Maternal effects are predicted to adaptively modulate offspring phenotypes according to the local environment (Mousseau and Fox, 1998; Marshall and Uller, 2007; Yin et al., 2019; Sánchez-Tójar et al., 2020). Females may influence their offspring prenatally through, for example, incubation temperature (Hepp et al., 2006; Mitchell et al., 2015) and by providing nutrients (Roseboom et al., 2006; de Rooij et al., 2010) or other resources, such as antioxidants (Blount et al., 2002), immune factors (Blount et al., 2002; Saino et al., 2002) and hormones (Schwabl, 1993; McCormick, 1999; Dloniak et al., 2006; Uller et al., 2007; Dantzer et al., 2013). These prenatal conditions can subsequently affect offspring sex ratios (e.g. Carter et al., 2017), development (Hepp et al., 2006; Mitchell et al., 2015), physiology (Blount et al., 2002; Saino et al., 2002; Boots and Roberts, 2012), behaviour (Eising et al., 2006; Partecke and Schwabl, 2008) and ultimately offspring fitness (e.g. Ruuskanen et al., 2012). What mediates these effects has however remained elusive (e.g. Groothuis et al., 2019; Bebbington and Groothuis, 2021).

Especially hormones are known to profoundly affect offspring phenotypes. In oviparous species, the effects of yolk androgen hormones, especially testosterone, on phenotypic traits of developing offspring have been extensively studied (Gil, 2008). Yolk testosterone levels have been found to vary between clutches, which can partly be explained by between female differences such as female age (Pilz et al., 2003), perceived predator risk (Coslovsky et al., 2012) and personality (Ruuskanen et al., 2018), and by environmental conditions, such as parasite load (Tschirren et al., 2004), or social conditions, such as mate attractiveness (Remeš, 2011) and breeding density (Groothuis and Schwabl, 2002; Eising et al., 2008; Remeš, 2011; Bentz et al., 2013). Despite studies showing that testosterone and other yolk androgens have largely disappeared after five days of incubation (Kumar et al., 2019), yolk testosterone has profound effects on offspring physiology and behaviour. Increased yolk testosterone levels have been found to stimulate nestling growth (Schwabl, 1996; Navara et al., 2005, 2006; Müller et al., 2007), decrease social dependence and neophobia (Daisley et al., 2005), and alter breath rate (Bentz et al., 2013), food intake and begging rates (Schwabl, 1996; Eising and Groothuis, 2003).

The effects of increased yolk testosterone levels are not always in the same direction, as some studies have reported decreased or unaffected begging rates (Pilz et al., 2004; Boncoraglio et al., 2006) and reduced growth (Sockman and Schwabl, 2000; Rubolini et al., 2006). Moreover, yolk testosterone can be deposited in a sex-specific way (Müller et al., 2002) and is known to induce sex-specific effects (Tschirren, 2015). Yolk testosterone has, for example, been found to promote growth in male offspring, while female growth was unaffected (Tschirren, 2015) or reduced (Saino et al., 2006). Likely, developmental trajectories, and their associated costs and fitness returns, are differentially affected in male and female offspring (Gil, 2008). This suggests a pivotal role for yolk testosterone in explaining how the sexes trade-off developmental processes differently. However, the interpretation of such sex-specific effects might be improved if the underlying molecular mechanisms that allow for such sex-specific effects are taken into account (Gil, 2008; Groothuis and Schwabl, 2008). Yet little is known on the molecular mechanisms that are responsible for these phenotypic changes in developing offspring caused by variation in yolk testosterone.

Epigenetic mechanisms have been suggested to facilitate effects of maternal androgens (Groothuis and Schwabl, 2008), specifically DNA methylation (Sepers et al., 2019). DNA methylation, the addition of a methyl group to a DNA nucleotide, interferes with the binding of transcription factors to the DNA (Bird, 2002; Moore et al., 2013; Yin et al., 2017). DNA methylation usually suppresses gene expression (Bird, 2002; Goldberg et al., 2007; Moore et al., 2013), specifically if methylated cytosines in CpG context (CG dinucleotides) are located nearby the transcription start site of a gene (Bird, 2002; Goldberg et al., 2007; Li et al., 2011; Moore et al., 2013; Laine et al., 2016). Therefore, DNA methylation is generally expected to affect the expression of phenotypic traits (Law and Jacobsen, 2010). Prenatal maternal effects on DNA methylation have been found in the offspring of vertebrates such as mice (St-Cyr and McGowan, 2015) and humans (Tobi et al., 2009, 2014). Studies in wild avian species have experimentally shown (Hukkanen et al., 2023) or suggested (Bentz et al., 2016) that maternal androgens can impact offspring methylation. To our knowledge, there is only one genome-wide study on the effects of experimentally elevated yolk androgens (in this case testosterone) on DNA methylation. In this study on zebra finches (*Taeniopygia guttata*), differentially methylated regions between treated and untreated individuals were found in or near genes that were also differentially expressed in the hypothalamus and amygdala (Bentz et al., 2021). However, this might be specific to males as female zebra finches were not included in this study. Sex differences in DNA methylation are known to exist in other bird species (Teranishi et al., 2001; Natt et al., 2014) and sex-specific differences in erythrocyte DNA methylation levels were previously found in the great tit (Verhulst et al., 2016). Therefore, to what extent DNA methylation is an underlying mechanism for yolk testosterone mediated sex-specific maternal effects remains largely unknown.

Here, we experimentally manipulated yolk testosterone levels in a population of wild great tits (*P. major*) and assessed pre-fledging effects on biometric measures, behavioural traits, and genome-wide DNA methylation to test whether yolk testosterone-induced changes in DNA methylation may explain sex-specific phenotypic effects. We expected that testosterone positively affects body mass and tarsus length in male great tits, but not in females (Tschirren, 2015), explained by higher competitiveness in testosterone treated males, observed by a higher begging rate and higher food reception (Schwabl, 1996; Eising and Groothuis, 2003). We expected testosterone treated males to have a lower stress response as a result of receiving larger amounts of food (van Oers et al., 2015). If sex-specific effects of testosterone on behaviour and biometry are mediated by DNA methylation, we expected the transcription start site regions of genes involved in sexual dimorphism, hormone receptor expression, steroid hormone secretion, neuron development, morphology and growth to be differentially methylated between the testosterone treated group and the control group, but only in within-sex comparisons. Our results show that sex-specific DNA methylation changes associate with sex-specific effects on biometry and behaviour caused by experimentally induced elevated yolk testosterone levels. This therefore strongly suggests that DNA methylation is indeed a mediator of post-hatching sex-specific effects of elevated yolk testosterone on offspring phenotypes, but these effects do not always induce a more competitive phenotype pre-fledging.

## Material and methods

This study was conducted in 2020 in a long-term nest box population in the Westerheide estate, near Arnhem, the Netherlands (52°01′00N, 05°50′30E). All information regarding number of eggs, nestlings, clutches and broods is provided in Supplementary tables S1, S2 and S5.

### Experimental injection protocol

We manipulated yolk androgen levels following a procedure that has been successfully applied before (*e.g.* Tschirren *et al*., 2005; Ruuskanen and Laaksonen, 2010) (described in detail in Appendix S1-A). Briefly, clutches were alternately assigned to be in either the testosterone treated group or the control group. On the day the fifth egg was laid, egg yolks in clutches assigned to the testosterone treated group were injected with 12 ng of testosterone dissolved in 5 µL of sesame oil, while egg yolks in control clutches were injected with 5 µL of sesame oil. Thereafter, injections were done each day for the newly laid egg. Clutch size, egg weight, the duration of the incubation period, and hatching success did not differ significantly between the treatment groups (Appendix S2-A). Although hatching success was low (58.4% for control broods and 61.6% for testosterone treated broods, Table S1), it was high compared to other injection studies (50-55%, e.g. Podlas et al., 2013; Ruuskanen et al., 2016; Bentz et al., 2021).

### Cross-fostering

To minimise the differences in rearing environment between nestlings from testosterone treated eggs and nestlings from control eggs, we used a partial cross-foster design as in Van Oers *et al*. (2015) and Sepers *et al*. (2021) (described in detail in Appendix S1-B). Briefly, we assigned broods to pairs on day two or three after hatching. Within a pair, the nestlings were weighed (to the nearest 0.01 g) and partially cross-fostered to create mixed broods containing nestlings from both treatments. Pairs of small broods (± three nestlings) were merged into one brood to lower the chance of desertion. We discarded 21 broods where the testosterone treated nestlings were one day younger than the control nestlings from all analyses (asynchronous pairs, Table S1), as this made it impossible to disentangle treatment effects from age effects. On day six after hatching, the nestlings were weighed and approximately 10 μL of blood was collected by brachial venipuncture. The samples were stored at room temperature in one mL of cell lysis buffer (Gentra Puregene Kit, Qiagen, USA) until further analysis for molecular sexing (following Griffiths et al., 1998) and to measure erythrocyte DNA methylation levels.

### Video recordings and analysis

Seven days after hatching, we installed an infrared spy-camera (Velleman, CMOS-camera, CAMCOLMBLAH2) to record nestling food solicitation behaviours. Cameras were connected to a digital video recorder placed outside the nest box (PV-500L2, LawMate International, Taipei, Taiwan). On day eight after hatching, we weighed nestlings and marked them with red acrylic paint to enable their identification on video recordings under infrared light. Subsequently, we recorded the brood for at least two hours between 07:00 and 15:00. One nest box was recorded between 15:30 and 17:30. At least 1.5 hours of video recordings were analysed using Adobe Premiere Pro 2021 (Adobe Inc.) by a single person who was blind with respect to nestling treatments. During each parental feeding visit, we scored (a) which nestlings showed a begging response right before feeding and (b) which nestling received food.

### Handling stress test

We conducted a handling stress test fourteen days after hatching, as described in Fucikova *et al*. (2009) and Sepers *et al*. (2023b), with two modifications. We measured handling stress for only one minute instead of two, and the nestlings were not socially isolated. Handling stress was measured by counting the number of breast movements (*i.e.* breath rate) during four subsequent bouts of each 15 seconds. Once all nestlings were tested separately, they were weighed and their tarsus length (callipers, ± 0.1 mm) was measured. Individual estimates for the handling stress response were obtained as described in Sepers *et al*. (2023b), using a linear mixed model (LMM) with the number of breaths per 15 second bout as dependent variable. The produced estimates (*i.e.* handling stress response) quantify the individual deviations from the average slope in breath rates over time and were extracted for further analysis.

### DNA methylation data generation and bioinformatics

From all blood samples collected on day six after hatching, we randomly selected four samples per brood of rearing (two samples from each treatment). In total, we selected 180 samples, of which 100 were from broods without an age difference between control and testosterone treated nestlings (Table S5). We assessed genome-wide DNA methylation levels using epiGBS2 (Gawehns et al., 2022). This is a reduced-representation DNA methylation laboratory protocol and a bioinformatics pipeline. Five libraries, each containing 36 barcoded samples, were prepared and sequenced as described in Gawehns *et al*. (2022) with the improvements reported in Sepers *et al*. (2023b) and described in detail in Appendix S3.

Raw reads were demultiplexed, checked, trimmed, filtered, merged, aligned and called for methylation using the reference branch of the epiGBS2 bioinformatics pipeline (Gawehns et al., 2022) with minor modifications. We aligned the reads to the *P. major* reference genome v1.1 (GCF_001522545.3) (Laine et al., 2016) and removed overlap between read pairs (Table S6). Methylation was called while ignoring the first four nucleotides in all reads and in CpG context only. Complementary CpG dinucleotides were merged using the R package *methylKit* v1.16.1 (Akalin et al., 2012). Subsequently, using a custom script, CpG sites (CpGs) with low coverage (< 10X) or very high coverage (> 99.9^th^ percentile), and CpGs that were not covered in at least 15 individuals in each treatment or with a high or low mean methylation level (<0.05 or >0.95 across all individuals) were discarded (Table S7), leading to a total of 169,215 CpGs in the final analysis.

### Statistical analysis

#### General

For all mixed models described below, unless described otherwise, we included treatment, sex and their interaction as fixed effects. Brood of origin and brood of rearing were included as random effects to account for non-independence of nestlings from the same brood.

We used a binomial error distribution and logit-link function for all generalized linear mixed models (GMMLs). The significance of the interaction of treatment with sex and/or the significance of treatment were determined with a Likelihood Ratio Test (LRT) by comparing a model with the interaction or factor of interest with a model without the interaction or factor of interest using the *anova* function in R.

All LMMs were run with ML estimation and we used backwards elimination of the interaction between treatment and sex based on the p-values provided by a type III analysis of variance via Satterthwaite’s degrees of freedom method using the *anova* function in R. In case the interaction was non-significant (p < 0.05 for biometry and behavioural data, p < 0.1 for DNA methylation data) it was deleted. The minimal adequate model always included treatment and sex. The analyses were done using the packages *lme4* v1.1.28 and *lmerTest* v3.1.3. Post-hoc comparisons were performed with the *lsmeans* function in the package *emmeans* v1.7.2. P-values were provided via the Satterthwaite’s degrees of freedom method and corrected for multiple testing with a Bonferroni correction.

All behaviour and biometry analyses were done in Rstudio v2021.9.2, while the analyses of the methylation calls were done with Rstudio v1.4.1717.

#### Statistical analysis biometry and behaviour

We used separate LMMs to analyse the effect of the treatment on weight on day two, day six, day eight and day 14 after hatching and on tarsus length and handling stress on day 14. Brood of rearing was not included when analysing the effect on weight on day two, as weighing happened before cross-fostering.

We analysed the effect of the treatment on the probability of begging (yes/no) and probability of getting fed at each parental visit using GLMMs. Nestling ID was included as random effect in addition to the fixed and random effects described in *General*.

#### Statistical analysis DNA methylation

To analyse the effect of the treatment on DNA methylation level per CpG, we used a GLMM in which the dependent variable was modelled as the fraction of the number of methylated Cs over the total number of analysed reads (*i.e.* coverage: number methylated of Cs plus unmethylated Cs per CpG) with the *cbind* function. This model was run for each CpG separately and included the fixed and random effects described in *General*. CpGs with a False Discovery Rate (FDR) (Benjamini and Hochberg, 1995) corrected p-value below 0.1 were considered as significantly differentially methylated CpGs. CpGs for which the effect of the treatment depended on sex were referred to as sex-specific differentially methylated sites (sex-specific DMS). We excluded models of CpGs that produced warnings other than singularity warnings. Furthermore, we corrected for potential overdispersion by excluding CpGs that fell out of the 95% Highest Density Interval (HDI) for the distribution of the dispersion statistic (Zuur et al., 2013) using the R package *HDInterval* v0.2.2 (Table S7).

Subsequently, each sex-specific DMS was assigned to a category. A significant (FDR corrected) difference between the treatments in both the females and males, while the within sex differences were in opposite directions, corresponded to the category “antagonistic DMS”. A significant difference between control and testosterone treated females corresponded to the category “female-specific DMS”, while a significant difference between control and testosterone treated males corresponded to the category “male-specific DMS”. No significant difference between the treatments in either the males or females, but some other significant difference corresponded to the category “other DMS” (*e.g.* significant difference between testosterone treated females and control males).

#### Gene annotation and ontology analyses

CpGs were assigned to the genomic regions as described in Sepers *et al*. (2021) and (2023a) using custom R scripts, R packages *GenomicFeatures* v1.42.3 (Lawrence et al., 2013) and *rtracklayer* v1.50.0 (Lawrence et al., 2009) and the *P. major* reference genome build v1.1, annotation version 102 (Laine et al., 2016). To facilitate interpretation of genes associated with these sex-specific DMS, we searched the literature for relevant studies on phenotypic effects, DNA methylation and gene expression. We specifically focused on genes with DMS within regulatory regions (promoter and TSS regions) and (groups of) genes with multiple DMS (Tables S17-S20). In addition, we identified enriched Gene Ontology (GO) terms using the ClueGO v2.5.8 (Bindea et al., 2009) plug-in for Cytoscape v3.9.1 (Shannon et al., 2003) following Sepers *et al*. (2024). The target lists consisted of all genes associated with either (1) antagonistic DMS, (2) female-specific DMS, or (3) male-specific DMS. The background list consisted of all genes associated with any of the analysed CpGs, except for 3,603 functionally uncharacterised genes (*i.e.* LOC genes). All ontologies were updated on 09-03-2022. We used Revigo (Supek et al., 2011) to summarise GO terms and eliminate redundant terms. We applied a semantic similarity cut-off value of 0.5 *sim_Rel_* (Schlicker et al., 2006).

## Results

### Biometry and behaviour

Nestlings hatching from testosterone injected eggs did not differ in weight from those hatching from control eggs on day two (treatment, LMM: F_1,34.10_ = 0.30, p = 0.59; Table S8), day six (treatment, LMM: F_1,33.19_ = 0.13, p = 0.72; Table S9) or day eight after hatching (treatment, LMM: F_1,27.43_ = 0.26, p = 0.61; Table S10). Sexes significantly differed in weight on day six after hatching (sex, LMM: F_1,137.06_ = 5.43, p = 0.02), but not on the other days (all p ≥ 0.25). The treatment effect on variation in weight on day two, six or eight did not differ between the sexes (treatment x sex, all p ≥ 0.22).

Testosterone treatment affected chick weights on day 14 after hatching in a sex-specific way (treatment × sex, LMM: F_1,110.24_ = 5.24, p = 0.02; Table S13). Testosterone treated females weighed significantly less than testosterone treated males (estimate ± SE = −0.98 ± 0.17, t-ratio_114.5_ = −5.64, p < 0.001; Figure 2), while the sexes in the control group did not differ significantly in weight (−0.41 ± 0.18, t-ratio_114.2_ = −2.28, p = 0.15), and treatment effects within sexes were non-significant (females: 0.48 ± 0.27, t-ratio_32.2_ = 1.74, p = 0.55, males: −0.09 ± 0.27, t-ratio_30.3_ = −0.33, p = 1.00).

Testosterone treated nestlings (probability ± SE: 0.50 ± 0.03) begged significantly less than control nestlings (0.57 ± 0.03) did (treatment, GLMM: *χ*^2^_1_ = 3.99, p < 0.05; Figure 1; Table S11). This did not result in different probabilities of being fed (treatment, GLMM: *χ*^2^_1_ = 1.02, p = 0.31; Table S12). The treatment effect on the probability of begging or getting fed did not differ between the sexes (treatment x sex, all p ≥ 0.18).

**Figure 1.**
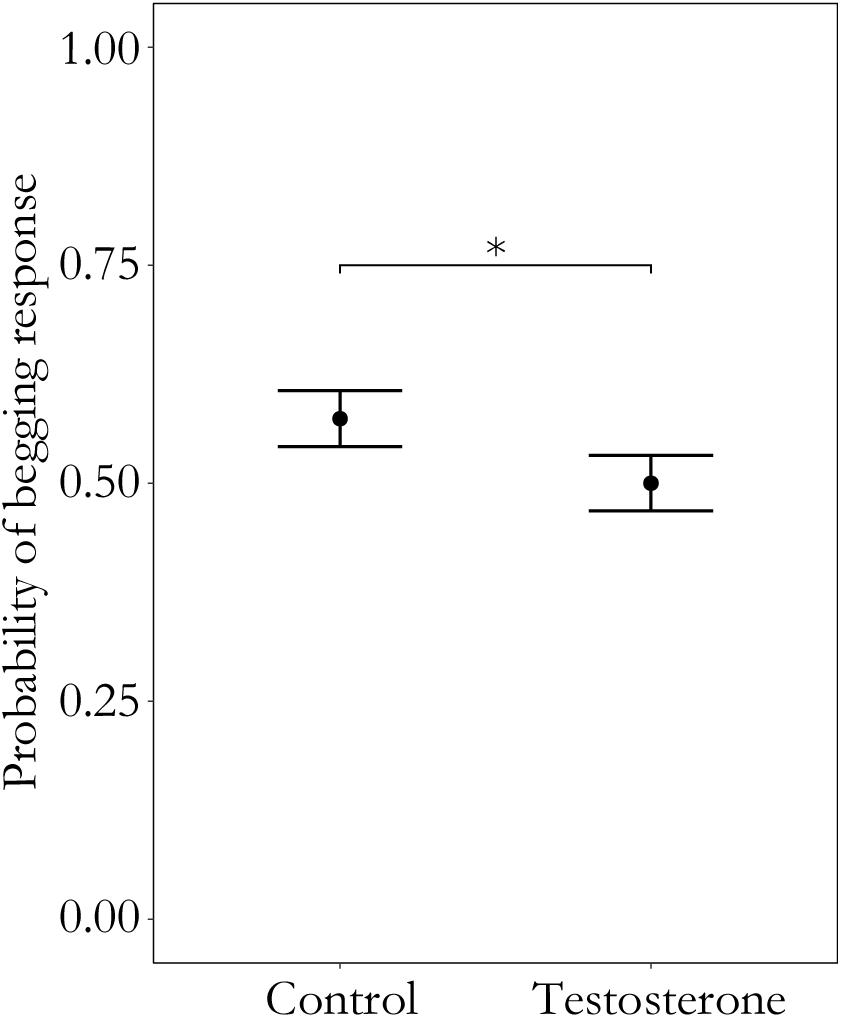
Begging probability. The effect of treatment on the average probability (± SE) of begging during parental visits. P-value below 0.05 but above 0.01 is indicated with *.

**Figure 2.**
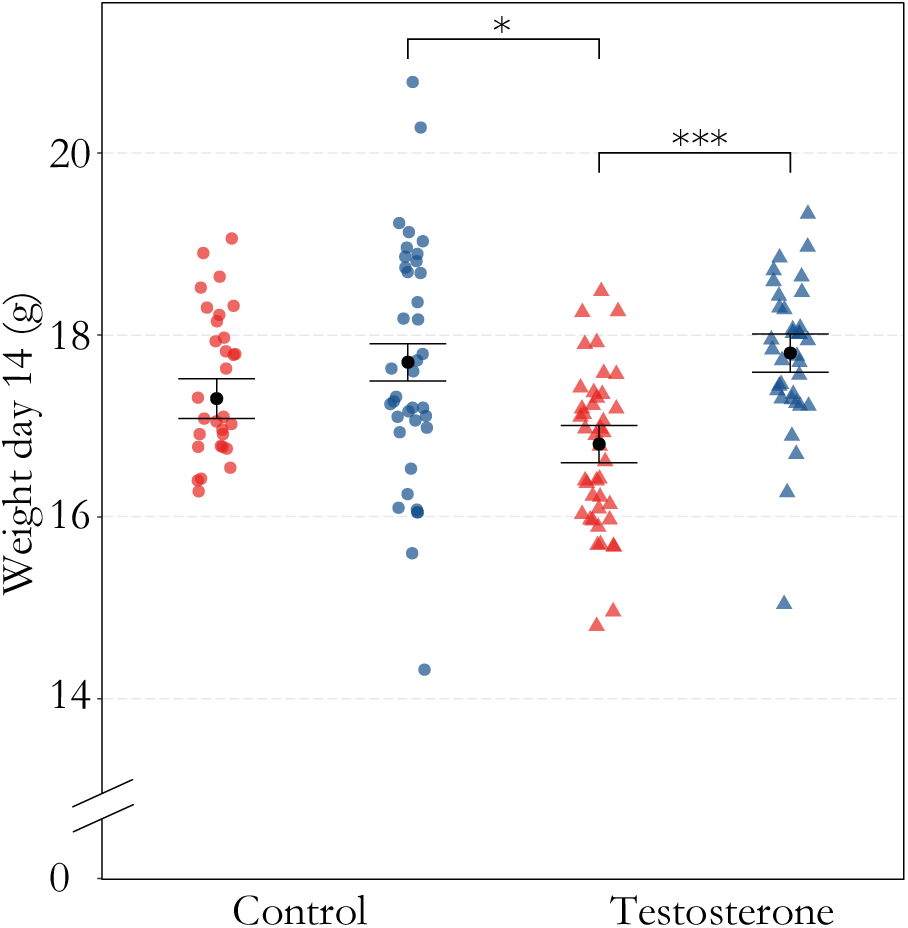
Weight day 14. Weight (g) on day fourteen after hatching for both sexes and control ● and testosterone ▴ treated nestlings. Red dots and triangles 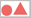 represent raw data points for females. Blue dots and triangles 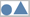 represent raw data point for males. Black dots represent the predicted marginal means for both sexes in both treatments. Error bars represent standard error of predicted marginal means. P-values below 0.05 but above 0.01 are indicated with *, p-values below 0.001 are indicated with ***.

We found a non-significant trend (treatment, LMM: F_1,27.54_ = 3.08, p = 0.09; Table S14) that testosterone treated individuals (mean ± SE = 19.30 ± 0.07) had longer tarsi on day 14 after hatching compared to control individuals (19.10 ± 0.07), after correcting for sex differences (sex, LMM: F_1,127.72_ = 61.30, p < 0.001). This tendency existed irrespective of the offspring sex (treatment × sex, LMM: F_1,120.93_ = 0.40, p = 0.53). Sex differences in the handling stress response (sex, LMM: F_1,124.63_ = 10.38, p = 0.002; Table S15) were not affected by the treatment (treatment × sex, LMM: F_1,122.21_ = 1.90, p = 0.17). Treatment itself also did not affect the handling stress response (treatment, LMM: F_1,32.85_ = 0.006, p = 0.94) when correcting for sex.

### Treatment effects on DNA methylation

While we did not find any DMS when comparing methylation between nestlings from testosterone treated eggs and control nestlings (all FDR corrected p-values > 0.1), we found 763 sites that showed a sex-specific treatment effect (Figure 3; Table 1). In 338 CpGs, testosterone significantly affected methylation in both sexes, but in different directions (antagonistic DMS). In 117 CpGs, we found a significant effect of testosterone on methylation in females, but not in males (female-specific DMS, Table 1), while the opposite pattern was found in 201 CpGs (male-specific DMS).

**Figure 3.**
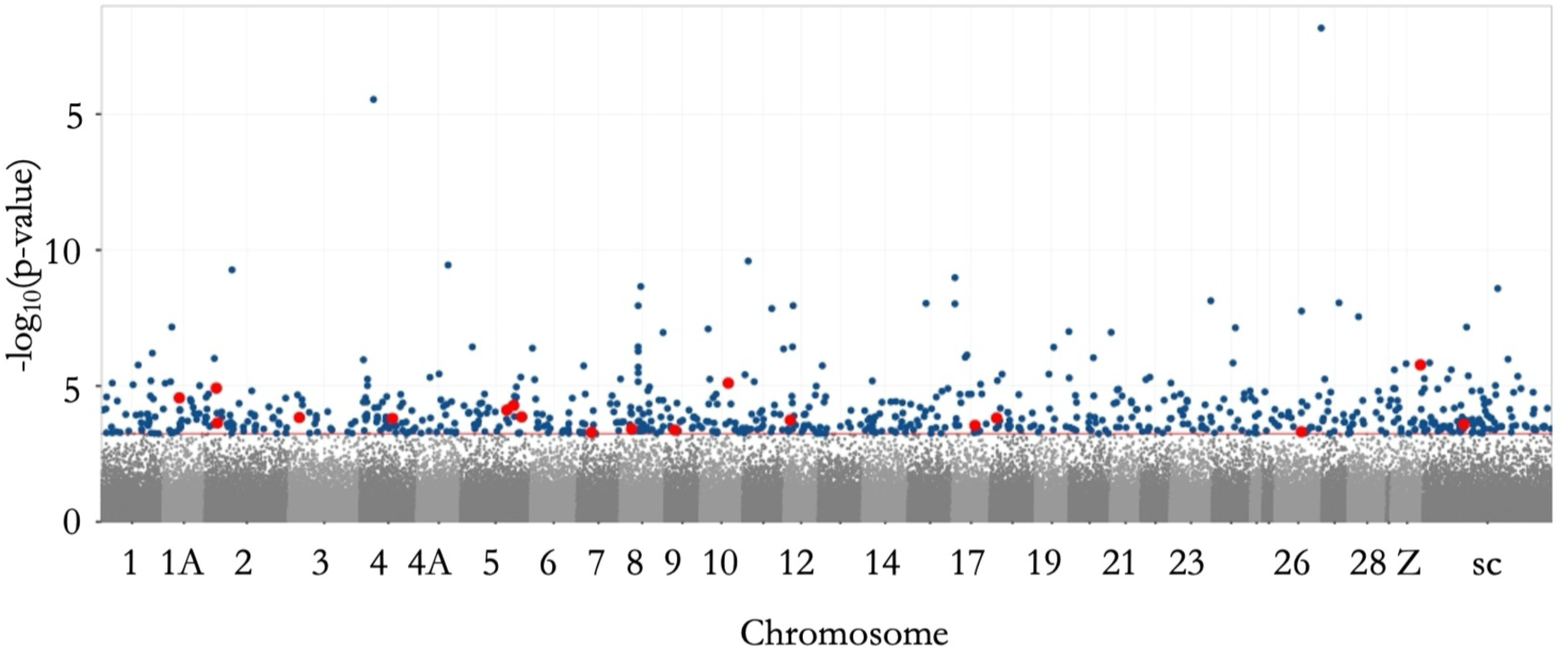
Manhattan plot of sex-specific DMS. Manhattan plot showing the significance of the interaction of treatment (control and testosterone) with sex in explaining variation in DNA methylation (−log_10_(p-values)). Each dot represents a CpG tested for a sex-specific treatment effect (169,215 CpGs). Dark blue dots represent CpGs with a significant sex-specific treatment effect. Red dots represent significant CpGs which are in the TSS region of a gene. The dotted red line marks the genome-wide significance threshold. The sites are plotted against the location of the associated site within the genome. Alternating colours help to differentiate adjacently displayed chromosomes. ChrZ is a sex chromosome, all the other chromosomes are autosomes. All unplaced scaffolds are merged into the category scaffolds (sc).

**Table 1.**
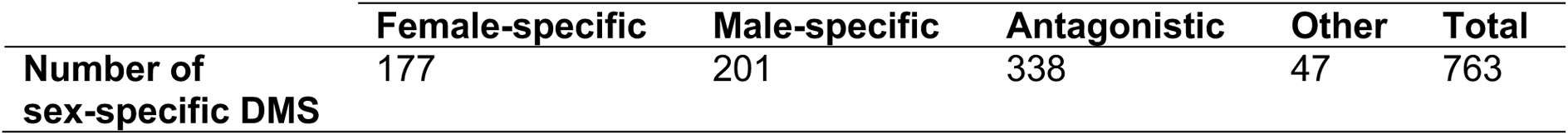
Number of sex-specific DMS assigned to different categories. The categories are based on whether the within sex differences between testosterone and control individuals are significant and the direction of the difference. Female-specific DMS: FDR corrected p-value < 0.1 between control and testosterone treated females only. Male-specific DMS: FDR corrected p-value < 0.1 between control and testosterone treated males only. Antagonistic DMS: a significant difference between the treatments in both sexes, while the within sex differences are in opposite directions. Other DMS: no significant difference between the treatments in either the males or females.

Out of these 763 CpGs, the 650 that could be annotated were found in or adjacent to 576 different genes, including 456 characterised genes. Out of the 650 annotated sex-specific DMS, 138 were found in the promoter region of a gene, including 18 situated in the TSS region. Furthermore, 392 sites were found in a gene body and 120 sites were found upstream or downstream of genes (Table S16).

Using the genes associated with antagonistic DMS, we detected 49 significantly enriched GO terms (FDR corrected p < 0.05; Tables S21 and S22), which clustered into 18 enrichments. The gene ontologies were mainly involved in nervous system development, cell adhesion, protein secretion, regulation of signaling pathways and transcription coregulator binding (Figure 4a).

**Figure 4.**
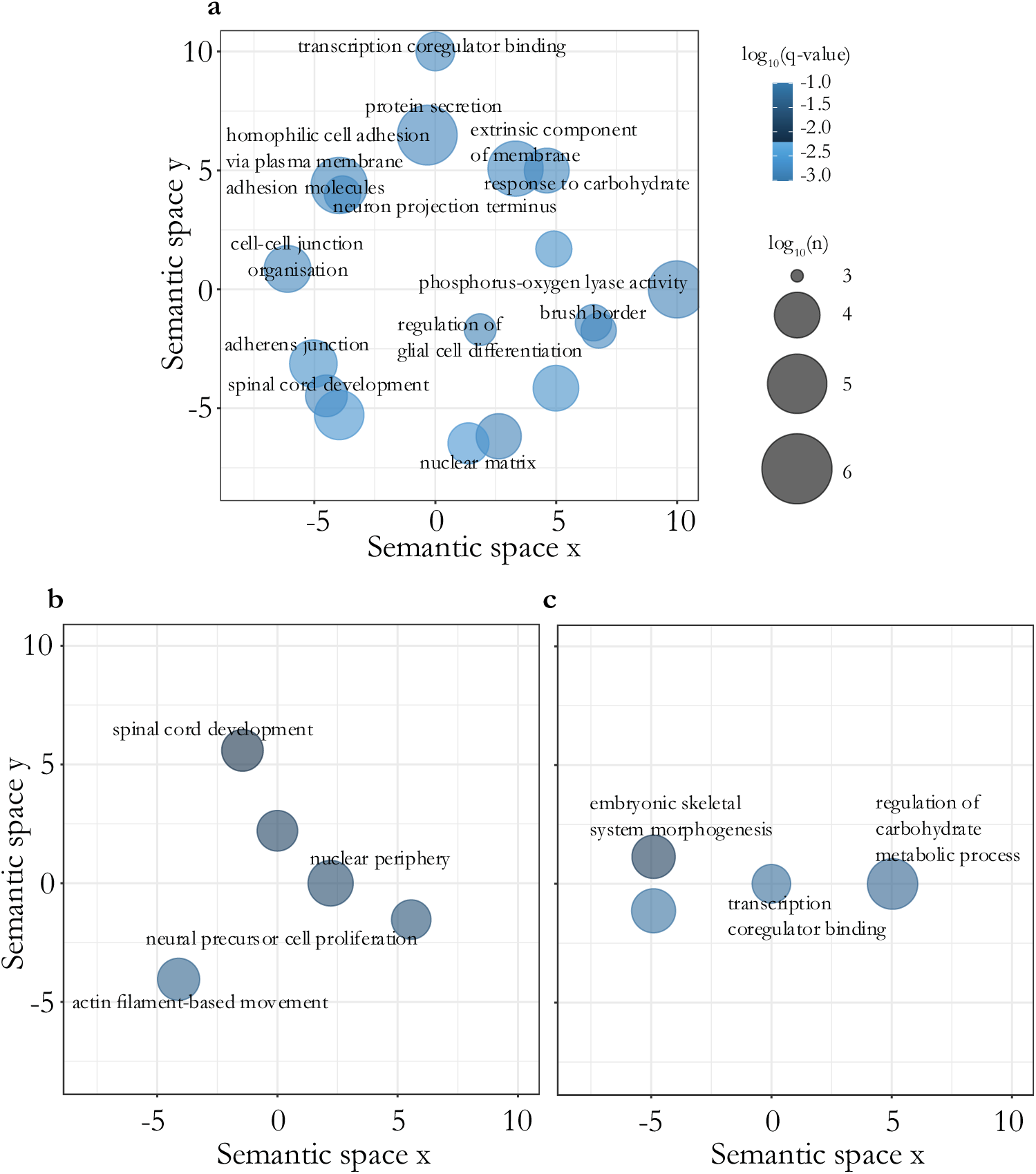
Representation of functional role of genes associated with sex-specific DMS. GO terms associated with sex-specific DMS for the categories (**a**) antagonistic DMS, (**b**) female-specific DMS and (**c**) male-specific DMS. The GO terms from ClueGO were merged based on semantic similarity using REVIGO. Colour scale indicates the FDR corrected p-value (q) (log_10_ scale) of the representative GO term for each merged category. Circle size shows the number of annotations (log_10_ scale) for the representative GO term in the underlying database. Larger circles represent more general terms, smaller circles represent more specific terms. GO terms were based on the ontologies biological process, molecular function and cellular component.

We detected nine significantly enriched GO terms (FDR corrected p < 0.05) when we conducted a GO analysis on genes associated with female-specific DMS only (Table S23). Five enrichments remained after merging. The gene ontologies were nuclear periphery, nuclear matrix, spinal cord development, neural precursor cell proliferation and actin filament-based movement (Figure 4b).

We also detected nine significantly enriched GO terms (FDR corrected p < 0.05) when we conducted a GO analysis on genes associated with male-specific DMS only (Table S23), which merged into three enrichments. The gene ontologies were transcription coregulator binding, embryonic skeletal system morphogenesis and regulation of carbohydrate metabolic process (Figure 4c).

## Discussion

Hormonally mediated maternal effects have profound effects on offspring phenotypes, often in a sex-specific way. DNA methylation is predicted to mediate these effects. The aim of this study was, therefore, to assess whether DNA methylation can potentially mediate sex-specific effects of experimentally manipulated yolk testosterone levels on postnatal biometry and behaviour. We found that experimentally elevated levels of yolk testosterone indeed affected individuals in a sex-specific way. Increased levels of yolk testosterone decreased the probability of begging and increased fledging mass differences between the two sexes. Experimental exposure to high levels of yolk testosterone also affected DNA methylation at 763 CpGs. Interestingly, the direction of this effect always depended on offspring sex.

We found a similar number of male-specific DMS and female-specific DMS, suggesting that the sexes are equally sensitive to an increase in testosterone in terms of DNA methylation, but at different CpG sites. As DNA methylation can affect gene expression, it is generally expected to affect the expression of phenotypic traits (Law and Jacobsen, 2010). Indeed, the biological functions of these genes were linked to phenotypic effects of experimentally elevated prenatal testosterone found in this study and in previous ones.

### Nervous system development and behaviour

Female-specific and antagonistic DMS were found in or near genes involved in nervous system development and behaviour (Table S17), which indicates a sex-specific effect of prenatal testosterone on brain organisation and behaviour. Indeed, experimentally increased yolk testosterone affected the brain anatomy differently in male and female chicks (*Gallus domesticus*) (Schwarz and Rogers, 1992). This was not reflected in the behavioural traits measured in our study, as we found no effect of exposure to high yolk testosterone on the handling stress response and it decreased the probability of begging, but regardless of sex. Although most studies reported stimulating effects of experimentally increased yolk testosterone on begging behaviour (Schwabl, 1996; Eising and Groothuis, 2003), decreased begging rates have been found before as well (Pilz et al., 2004). As the treatment did not affect nestling mass on day two, six and eight after hatching, it is unlikely that a difference in nestling quality or the need of food explains the treatment effect on begging behaviour. It is more likely that this was facilitated by structural differences, possibly mediated by DNA methylation.

### Mass, growth and metabolism

Exposure to high yolk testosterone increased fledging mass differences between the two sexes, since it had a negative effect on female but not on male fledging mass. However, this result must be interpreted with caution as the treatment only caused a marginally significant difference in fledging mass between females. This antagonistic effect of experimentally elevated yolk testosterone on growth has also been found in other bird species (Saino et al., 2006; Holmes and Schwabl, 2022). Sex-specific effects on fledging mass are also supported by the functions of several of the genes in which male-specific and antagonistic DMS were found (Table S18).

As described before, the decreased probability of begging of testosterone treated nestlings did not lead to a decreased nestling weight at earlier stages, possibly because sibling competition was low, which is supported by our results that they were not fed less. A likely explanation for the late appearance of an effect on weight is that when the nestlings from the testosterone group remain to beg less often than those from the control group when nestling competition increases, the female offspring may suffer the most in such suboptimal conditions (de Kogel, 1997; Martins, 2004). Indeed, several studies suggest that male great tit nestlings have a competitive advantage in suboptimal conditions (Dhondt, 1970; Drent, 1984; Smith et al., 1989; Lessells et al., 1996; Oddie, 2000).

### Reproduction

Several of the male-specific and antagonistic DMS were associated with male fertility (Table S19), which is confirmed by experimentally elevated yolk testosterone effects on fecundity and other traits that are important for reproduction, such as plumage development and sexual display (Eising et al., 2006; Partecke and Schwabl, 2008; Galván and Alonso-Alvarez, 2010), testis and egg size (Uller et al., 2005), and egg fertility and laying activity (Rubolini et al., 2007). This strongly suggests an effect of exposure to high yolk testosterone on fecundity and ultimately fitness via DNA methylation. To confirm this, it would be particularly interesting to assess DNA methylation, sperm quality and fitness measures, such as lifetime reproductive success, during subsequent breeding seasons.

### Oxidative stress

We found several sex-specific DMS in genes associated with DNA binding, oxidative stress and DNA repair (Table S20). Studies on the phenotypic level support our findings on methylation differences of genes related to oxidative stress and DNA repair, as experimentally increased yolk testosterone increases metabolic rate in nestlings and adults (Tobler et al., 2007; Ruuskanen et al., 2013), which might cause more oxidative damage (Feuerbacher and Prinzinger, 1981; Buchanan et al., 2001). Indeed, experimentally increased yolk testosterone has been found to reduce DNA damage repair efficiency (Treidel et al., 2013) and increase telomere shortening (Parolini et al., 2019). Furthermore, experimentally increased yolk testosterone reduced plasma antioxidant levels in male, but not in female nestlings (Tobler and Sandell, 2009), suggesting differences in the degree of oxidative stress or differences in investment in DNA repair or protection between the sexes. Overall, regulation of genes related to oxidative stress and DNA repair via DNA methylation likely explains sex-specific effects of elevated prenatal yolk testosterone on oxidative stress.

### Future avenue

Sex differences in DNA methylation are known to exist and have been confirmed in humans (Liu et al., 2010; McCarthy et al., 2014; Yousefi et al., 2015; Gatev et al., 2021; Solomon et al., 2022) and chickens (Teranishi et al., 2001; Natt et al., 2014). The mechanisms that account for sex-specific methylation patterns remain elusive. These differences might be driven by hormones, which could be confirmed with the effects of increased levels of yolk testosterone on DNA methylation in this study. However, we cannot exclude the possibility that genetic variation plays a role too. DNA methylation is not only influenced by environmental factors (Sepers et al., 2021, 2024), but also by genetic variation (Höglund et al., 2020; Sepers et al., 2023a) or by genotype by environment interactions (Sepers et al., 2019). As the treatment effect on DNA methylation depended on offspring sex, and because the sex chromosomes are highly differentiated in carinate birds (see Solari, 1993; Graves, 2014), we expect that the epigenetic response to enhanced yolk testosterone will be genotype-dependent for at least part of the CpGs. In birds, sex is genetically determined with females being the heterogametic sex (ZW) and males being the homogametic sex (ZZ). None of the DMS was located on the Z chromosome, which is currently the only available sex chromosome in the reference genome, but as genetic variation can affect methylation of distant CpGs (Höglund et al., 2020; Sepers et al., 2023a), genetic variation on the sex chromosomes can potentially induce sex-specific methylation patterns on the autosomes. To be able to separate maternal effects from sex-specific genetic effects on DNA methylation, quantitative trait loci should be mapped for the sex-specific DMS.

## Conclusion

Our study suggests that in contrast to our expectations, experimentally enhanced levels of yolk testosterone resulted in a less competitive phenotype, especially in females. Testosterone affected DNA methylation in genes belonging to different pathways for males and females, pointing to a difference in costs and benefits of being exposed to higher yolk testosterone between the sexes. Hence, mothers may trade-off investment in males and females by varying testosterone deposition in their clutch. Differential methylation of genes involved in metabolism, mass and development align with the phenotypic effects of elevated yolk testosterone, suggesting that DNA methylation induced by elevated yolk testosterone can affect nestling mass and behaviour. In conclusion, our results support the hypothesis that DNA methylation variation caused by maternal hormones deposited in the egg can be a mediator for sex-specific effects during early development.

## Author contributions

KO, SR and BS conceived the study. BS conducted the experiment and collected all samples and biometry and handling stress data. BS and TM collected the video recordings and analysed the corresponding data. BS conducted the remaining bioinformatic and statistical analyses. CM conducted the laboratory work. BS, SR and KO drafted the manuscript. KO supervised the study. All authors contributed to editing the manuscript.

## Supporting information

Sepers_et_al_Supplement

## Acknowledgements

We would like to thank Youri van der Horst, Lian Grabijn and Renske Kriesels for fieldwork assistance, Imke Broekhuis for scoring the video footage and Martijn van der Sluijs for laboratory work. We are grateful to Geldersch Landschap & Kasteelen for the permission to conduct fieldwork in Westerheide.

## Data and availability statement

The raw genomic datasets are deposited on NCBI under BioProject PRJNA208335 under the SRA accessions SRX22027777, SRX22027778, SRX22027779, SRX22027780 and SRX22027781. Data related to brood characteristics and biometry are archived in the SPI-Birds Database (https://spibirds.org/en; Culina et al., 2021) and can be requested. The epiGBS2 bioinformatics pipeline can be accessed on Github (https://github.com/nioo-knaw/epiGBS2). All other data sets and scripts are deposited in the Dryad repository and will be made publicly available upon acceptance.

## Ethics approval statement

All animal experiments involved in this study were reviewed and approved by the Institutional Animal Care and Use Committee (NIOO-IvD) and were licenced by the CCD (Central Authority for Scientific Procedures on Animals; AVD-801002017831) to KO.

