## Supplementary material for "DNA methylation associates with sex-specific effects of experimentally increased yolk testosterone in wild nestlings": Sepers_et_al_Supplement

**Table of contents**

| | Page |
|---|---|
| **Appendix S1: Methods** | **2-5** |
| Table S1: Overview of number of eggs, nestlings and broods | 2 |
| Table S2: Overview of number of nestlings and broods on day 2, 6, 8 and 14 | 2 |
| A. Detailed description experimental injection protocol | 3 |
| B. Detailed description of the cross-fostering procedure | 4 |
| References Appendix S1 | 5 |
| **Appendix S2: Clutch and brood characteristics** | **6** |
| Table S3: Model pre-injection egg weight | 6 |
| Table S4: Model hatching success | 6 |
| **Appendix S3: epiGBS2 laboratory protocol** | **7** |
| **Appendix S4: Bioinformatics and statistical analyses DNA methylation** | **8** |
| Table S5: Overview number of nestlings and broods DNA methylation analysis | 8 |
| Table S6: Average number of reads per library and individual | 8 |
| Table S7: Number of CpGs | 8 |
| **Appendix S5: Full statistical models biometry and behaviour** | **9-11** |
| Table S8: Model weight day two | 9 |
| Table S9: Model weight day six | 9 |
| Table S10: Model weight day eight | 9 |
| Table S11: Model begging probability day eight | 10 |
| Table S12: Model probability of getting fed day eight | 10 |
| Table S13: Model weight day 14 | 10 |
| Table S14: Model tarsus length day 14 | 11 |
| Table S15: Model handling stress day 14 | 11 |
| **Appendix S6: Gene annotation and GO analysis** | **12-19** |
| Table S16: Number and annotation of sex-specific DMS | 12 |
| Table S17: Literature nervous system development and behaviour | 13 |
| Table S18: Literature growth and metabolism | 13 |
| Table S19: Literature male fertility | 13 |
| Table S20: Literature oxidative stress and DNA repair | 13 |
| References Tables S17-S20 | 14-15 |
| Table S21: Enriched GO terms antagonistic DMS | 16-17 |
| Table S22: Enriched GO terms antagonistic DMS | 17 |
| Table S23: Enriched GO terms female-specific and male-specific DMS | 18 |

**Appendix S1: Methods**

***Table S1.*** *Overview of the number of injected eggs, nestlings and broods included in the experiment. Number of eggs or nestlings and from how many broods of origin they originated from are provided for both treatments. If applicable, number of pairs of broods are also provided (either cross-fostered or merged into one brood of rearing). A pair consists of broods from opposite treatments. The nestlings in both broods are the same age (day two) in synchronous pairs, the control nestlings are one day older (day three) than the testosterone nestlings in asynchronous pairs.*

| | **Control** | | **Testosterone** | | **CF pairs** | **Merged pairs** | **Total pairs** |
|---|---|---|---|---|---|---|---|
| | **Eggs or nestlings** | **Broods origin** | **Eggs or nestlings** | **Broods origin** | | | |
| **Injected** | 311 | 37 | 308 | 37 | NA | NA | NA |
| **Predated/abandoned/ excluded*** | 32 | 1 | 50 | 5 | NA | NA | NA |
| **Incubation length known** | 275 | 33 | 258 | 30 | NA | NA | NA |
| **Not predated/ abandoned/excluded** | 279 | 36 | 258 | 32 | NA | NA | NA |
| **Unhatched** | 116 | 36 | 99 | 32 | NA | NA | NA |
| **Hatched** | 163 | 36 | 159 | 32 | NA | NA | NA |
| **Synchronous pair** | 79 | 17 | 87 | 17 | 12 | 5 | 17 |
| **Asynchronous pair** | 53 | 13 | 64 | 13 | 8 | 5 | 13 |

** Out of the experimental 74 clutches, four did not hatch at all because they got predated or were abandoned (one control clutch and three testosterone clutches). Furthermore, the nest box of one testosterone treated clutch was taken over by another great tit pair, creating a mixed brood, and one clutch contained two uninjected eggs, because the female laid two more eggs after incubation had started. This left us with 68 broods with 537 eggs, of which 279 eggs in 36 control broods and 258 eggs in 32 testosterone broods.*

***Table S2.*** *Overview of the number of nestlings and broods included in the experiment. Number of nestlings per sex and from how many broods of origin and broods of rearing they originated from are provided for both treatments on two, six, eight and 14 days after hatching. These numbers indicate how many individuals were included in the biometry, behavioural and survival analyses. The numbers go down over the season because some nestlings died.*

| | **Control** | | | | **Testosterone** | | | | **Broods** | **Broods** |
|---|---|---|---|---|---|---|---|---|---|---|
| | **♀** | **♂** | **?** | **Total** | **♀** | **♂** | **?** | **Total** | **origin** | **rearing** |
| **Asynchronous day two/three*** | 24 | 27 | 2 | 53 | 25 | 33 | 6 | 64 | 26 | 21 |
| **Synchronous day two** | 36 | 41 | 2 | 79 | 45 | 41 | 1 | 87 | 34 | 29 |
| **Pre-fledging, only synchronous pairs:** | | | | | | | | | | |
| **Day six** | 36 | 41 | NA | 77 | 45 | 41 | 1 | 87 | 34 | 29 |
| **Day eight** | 32 | 38 | NA | 70 | 43 | 36 | NA | 79 | 34 | 27 |
| **Recording day eight** | 19 | 23 | NA | 42 | 25 | 23 | NA | 48 | 25 | 16** |
| **Day 14** | 30 | 38 | NA | 68 | 42 | 35 | NA | 77 | 34 | 27 |

** Excluded from the analyses.*
*** Due to technical malfunctions, we were able to analyse the recordings from 16 broods.*

### A. Detailed description experimental injection protocol

We manipulated yolk androgen levels following a procedure that has been successfully applied before (*e.g.* Tschirren *et al.*, 2005; Ruuskanen and Laaksonen, 2010). Clutches were assigned to be in either the control group or the testosterone treated group, *i.e.* we used a between-clutch experimental design. The treatment of the first injected clutch was randomly assigned after which the treatments were alternated between clutches. On the day the fifth egg was laid, all eggs were taken from the nest box and replaced by dummy eggs. As great tits usually do not start incubation before the last egg is laid (Haftorn, 1981; Álvarez and Barba, 2014) and the average clutch size in this population was 8.26 ± 1.61 (mean ± SD) in the year before, this ensured that the eggs were injected prior to incubation, mimicking maternal testosterone deposition as much as possible. We put the great tit eggs in a container containing cotton balls and took them to a field station in Westerheide. Eggs were weighed (digital scale, ± 0.01 g) and placed on cotton padding on a wooden platform. The yolk was detected by illuminating the egg from below with a LED light (PowerBeam schouwlamp, MS Broedmachines, EAN: 7436950762744) through a hole in the wooden platform. The shell was cleaned with a cotton swap and 70% ethanol and punctured with a disposable 25G needle (BD Microlance). Subsequently, 5 µL of sesame oil or 12 ng of testosterone dissolved in 5 µL sesame oil was slowly injected into the yolk using a 10 µl syringe (Hamilton, 701RN) with a 26G needle (Hamilton, HAMI7758-02). As we aimed to elevate yolk testosterone to concentrations within the population's natural range in the majority of eggs, while still inducing detectable effects on biometry (Tschirren, 2015), we injected 12 ng testosterone per egg (approximately twice the standard deviation of yolk testosterone concentrations in great tit egg from a nearby study population: mean ± SD = 22.9 ± 6.2 ng/yolk in Ruuskanen et al., 2016). After injection, the syringe was slowly removed and the shell was cleaned with a cotton swap and 70% ethanol. Subsequently, the hole was sealed with tissue adhesive (3M Vetbond). The injected eggs were marked with a pencil and returned to the nest box of origin, after which the dummy eggs were taken out.

After injecting the first five eggs, clutches were checked daily to inject newly laid eggs following the same protocol until no new eggs were laid and incubation had started. If incubation had started, a heat pack (Rubytec Saba handwarmer) was placed over the eggs to keep them warm during transport. The clutches were left undisturbed until two days before the estimated hatch date. From then onwards, they were checked daily to determine the exact date of hatching (day 0).

A separate syringe was used for each treatment. The needle was cleaned with 95% ethanol between injections. The syringes were cleaned in between clutches by rinsing them first with 70% ethanol and then with sesame oil. At the end of each day, the syringes were rinsed with ethanol and left to dry until the next day. The syringes were rinsed with sesame oil the next morning before injections started.

*Testosterone solution*

The testosterone solution consisted of 12 ng of testosterone dissolved in 5 µL sesame oil. The testosterone solution was prepared by dissolving 20 mg testosterone (Sigma-Aldrich, SKU: 86500, ≥ 99% hormone) in 20 mL ethanol in a sterile 50 mL tube. Next, to create a batch solution, 19.2 µL of the highly concentrated solution was taken and mixed with 8 mL sesame oil (*Sesamum indicum,* Sigma-Aldrich*,* 85067) after which all the ethanol evaporated. Next, the solution was divided over polypropylene vials (Waters, SKU: 186002639) for daily use (200 µL of the batch solution each). Other vials were filled with just 200 µL of sesame oil, those were used for the eggs in the control group. The solutions were prepared as sterile as possible by pipetting in a flow chamber and autoclaving the vials and sesame oil before use (20 minutes at 120 °C). The vials were stored in darkness at room temperature. During injection, vials were replaced daily to minimise contamination.

### B. Detailed description of the cross-fostering procedure

We assigned broods to pairs on day two or three after hatching. From the 68 broods, we were able to create 30 pairs of broods from different treatments. We made two types of pairs. Either, a pair consisted of two broods that had different treatments and zero days difference in hatching date (all day two, 17 synchronous pairs; 166 nestlings from 34 original broods) or the control nestlings were one day older compared to the testosterone treated nestlings (13 asynchronous pairs, 117 nestlings from 26 original broods; Table S1).

Within a pair, the nestlings were partially cross-fostered to create mixed broods of both testosterone treated and control nestlings. Nestlings within broods were weighed and ranked based on their weight. We alternated ranking between pairs, so in one pair the heaviest nestling in each brood got the highest rank, while in a different pair the lightest nestling in each brood got the highest rank. This way, we avoided a bias in weight between cross-fostered (moved) and uncross-fostered (unmoved) nestlings. Subsequently, one brood received all the odd ranked nestlings, while the other brood received all the even ranked nestlings. This was decided randomly by flipping a coin. This way, each brood consisted of about as many testosterone treated as control nestlings while the original brood size was unaffected and weight differences between moved and unmoved nestlings were minimised (*i.e.* minimal differences between both treatments in the same brood) (van Oers et al., 2015).

If broods were small (± three nestlings), the nestlings were put into one brood (*i.e.* merged) instead of cross-fostered to lower the change of abandonment. The parent pair that received all nestlings was alternated between treatments in order to balance the fact that the treatments were evenly distributed to being either moved or unmoved.

This resulted in 50 broods of rearing (20 cross-fostered pairs and 10 merged pairs; Table S1). In 21 broods of rearing, the testosterone treated nestlings were one day younger than the control nestlings (asynchronous pairs). As this made it impossible to disentangle treatment effects from age effects, we excluded those 21 broods of rearing from the analyses on the nestling data.

To be able to recognise individual nestlings, we selectively plucked down on the head and back before cross-fostering. These codes were visible until day six after hatching, when each nestling was ringed with a uniquely numbered aluminium band (Vogeltrekstation, the Netherlands), and weighed (to the nearest 0.01 g).

**References Appendix S1**

Álvarez, E., and Barba, E. (2014). Incubation and hatching periods in a Mediterranean Great Tit *Parus major* population. *Bird Study* 61, 152–161. doi: 10.1080/00063657.2014.908819

Haftorn, S. (1981). Incubation during the Egg-Laying Period in Relation to Clutch-Size and Other Aspects of Reproduction in the Great Tit Parus major. *Ornis Scandinavica (Scandinavian Journal of Ornithology)* 12, 169–185. doi: 10.2307/3676074

Ruuskanen, S., Darras, V. M., Vries, B. de, Visser, M. E., and Groothuis, T. G. G. (2016). Experimental manipulation of food availability leads to short-term intra-clutch adjustment in egg mass but not in yolk androgen or thyroid hormones. *Journal of Avian Biology* 47, 36–46. doi: 10.1111/jav.00728

Ruuskanen, S., and Laaksonen, T. (2010). Yolk hormones have sex-specific long-term effects on behavior in the pied flycatcher (Ficedula hypoleuca). *Hormones and Behavior* 57, 119–127. doi: 10.1016/j.yhbeh.2009.09.017

Tschirren, B. (2015). Differential Effects of Maternal Yolk Androgens on Male and Female Offspring: A Role for Sex-Specific Selection? *PLOS ONE* 10, e0133673. doi: 10.1371/journal.pone.0133673

Tschirren, B., Saladin, V., Fitze, P. S., Schwabl, H., and Richner, H. (2005). Maternal yolk testosterone does not modulate parasite susceptibility or immune function in great tit nestlings. *Journal of Animal Ecology* 74, 675–682. doi: 10.1111/j.1365-2656.2005.00963.x

van Oers, K., Kohn, G. M., Hinde, C. A., and Naguib, M. (2015). Parental food provisioning is related to nestling stress response in wild great tit nestlings: implications for the development of personality. *Front. Zool.* 12, 10. doi: 10.1186/1742-9994-12-s1-s10

### Appendix S2: Clutch and brood characteristics

The pre-injection egg weight did not differ between control and testosterone treated broods (treatment, LMM: $F_{1,73.78}$ = 1.09, p = 0.30; Table S3).

Testosterone treated broods (mean ± SE = 8.66 ± 0.22 eggs) and control broods (8.50 ± 0.23 eggs) also had similar clutch sizes (Wilcoxon rank sum test; treatment; W= 532, p = 0.58).

From 63 broods (33 control and 30 testosterone treated broods) we knew exactly when incubation started (Table S1). Testosterone-treated broods (mean ± SE = 12.80 ± 0.16 days) were not incubated longer than control-treated broods (12.70 ± 0.13 days), showing that testosterone did not delay embryonic development (one-sided Wilcoxon rank sum test; treatment; W = 467, p = 0.34).

While the hatching success was low for both testosterone-treated broods (mean ± SE = 0.63 ± 0.03) and control broods (0.59 ± 0.03), there was no significant difference between the treatments (GLM; treatment; $\chi^2_{1,66}$ = 0.57; p = 0.45; Table S4).

***Table S3.*** *Model pre-injection egg weight. Table consists of all factors tested in the linear mixed model with maximum likelihood (ML) estimation was used. Egg weight before testosterone or control injection was included as the dependent variable. Treatment was included as a fixed effect and brood ID (i.e. brood of origin, var = 0.0094) as a random effect to control for non-independence of eggs from the same female. The estimate, degrees of freedom (numerator df and denominator df), the test statistic (F-value) and the significance (p-value) are given.*

| Egg weight | Estimate ± SE | Num. df | Denom. df | F-value | P-value |
|---|---|---|---|---|---|
| Intercept | 1.66 ± 0.02 | 1 | 73.73 | - | - |
| Treatment (T) | -0.02 ± 0.02 | 1 | 73.78 | 1.09 | 0.30 |

***Table S4.*** *Model hatching success. Table consists of all factors tested in the generalized linear model with binomial distribution and logit link. The dependent variable was the per brood fraction of the number of hatched eggs over the clutch size using the cbind function in R. Treatment was included as a fixed effect. The estimate, degrees of freedom (numerator df and denominator df), the test statistic ($\chi^2$ value) and the significance (p-value) are given.*

| Hatching success | Estimate ± SE | Num. df | Denom. df | $\chi^2$ | P-value |
|---|---|---|---|---|---|
| Intercept | 0.34 ± 0.12 | | | - | - |
| Treatment (T) | 0.13 ± 0.18 | 1 | 66 | 0.57 | 0.45 |

### Appendix S3: epiGBS2 laboratory protocol

Individuals belonging to the same pair were placed on the same library and pairs with matching hatch dates were divided over different libraries to avoid a correlation between library and hatch date. From each sample, 800 ng DNA was fragmented with *Msp*I (NEB) and large fragments were removed with beads (0.8x AMPure XP beads, Beckman Coulter). Next, we ligated the fragments to a unique adapter combination for identification of reads after sequencing. Subsequently, we pooled the fragments of all 36 samples and removed small fragments (NucleoSpin Gel & PCR Cleanup Kit, Macherey-Nagel). Next, the chemical sodium bisulfite was added to convert unmethylated cytosines (Cs) to thymines (Ts). Bisulfite-PCR amplification was conducted using the KAPA HIFI Uracil + hotstart ready mix (Roche) and 15 PCR cycles. Finally, the libraries were sequenced by Novogene (Novogene (HK) Company Limited, Hong Kong) on an Illumina HiSeq X (150 bp paired-end, directional reads). PhiX DNA (12%) was used to account for the depletion of cytosines in the bisulfite-treated data and allow for proper base calling calibration.

### Appendix S4: Bioinformatics and statistical analyses DNA methylation

***Table S5.*** *Overview of the number of nestlings and broods included in the DNA methylation analysis. Number of nestlings per sex and from how many broods of origin and broods of rearing they originated from are provided for both treatments. One sample per individual was included. These numbers indicate how many individuals were included in the pre-fledging DNA methylation analysis.*

| | Control | | | Testosterone | | | Broods origin | Broods rearing |
|---|---|---|---|---|---|---|---|---|
| | ♀ | ♂ | Total | ♀ | ♂ | Total | | |
| **Methylation synchronous** | 23 | 27 | 50 | 27 | 23 | 50 | 28 | 25 |
| **Methylation asynchronous*** | 18 | 22 | 40 | 18 | 22 | 40 | 26 | 20 |

* *excluded from the analyses*

***Table S6.*** *Average number of reads per library, average number of reads per individual after demultiplexing and filtering and average mapping efficiency per individual. Ranges are also provided.*

| | **Average (range)** |
|---|---|
| **# raw reads per library (R1+R2)** | 943,656,927 (93,572,4052 – 951,230,034) |
| **# filtered & demultiplexed reads per individual (R1+R2)** | 11,434,025 (1,355,424 – 36,679,004) |
| **mapping efficiency per individual** | 47.81% (43.10% - 51.40%) |

***Table S7.*** *Number of CpGs before and after filtering. Numbers are also provided for both models.*

| | **Number of unique CpGs** | |
|---|---|---|
| **Raw** | 4,304,738 | |
| **Destranding** | 2,373,607 | |
| **Coverage ≥ 10x** | 1,894,308 | |
| **≤ 99.9$^{th}$ percentile of coverage** | 1,894,305 | |
| **N ≥ 15** | 400,672 | |
| **0.05 ≥ mean meth ≤ 0.95** | 169,215 | |
| | **Treatment × sex** | **Treatment + sex** |
| **Input models** | 169,215 | 168,452 |
| **Output models** | 169,197 | - |
| **Removal of warnings** | 162,265 | - |
| **≤ 95% HDI dispersion statistic** | 157,039 | - |
| **FDR q-value < 0.1** | 763 | 0 |

### Appendix S5: Full statistical models biometry and behaviour

***Table S8.*** *Weight day two minimal adequate model. Table consists of all factors tested in the linear mixed model with the weight from two-day-old testosterone treated and control nestlings as the dependent variable. Brood of origin (var = 3614) was included as a random factor. The estimate, degrees of freedom (numerator df and denominator df), the test statistic (F-value) and the significance (p-value) are given.*

| Weight day two | Estimate ± SE | Num. df | Denom. df | F-value | P-value |
|---|---|---|---|---|---|
| **Minimal adequate model** | | | | | |
| Intercept | 2.96 ± 0.17 | 1 | 41.56 | - | - |
| Treatment (T) | 0.12 ± 0.23 | 1 | 34.10 | 0.30 | 0.59 |
| Sex (male) | 0.11 ± 0.09 | 1 | 136.10 | 1.36 | 0.25 |
| **Dropped terms** | | | | | |
| Treatment (T) × Sex (male) | 0.21 ± 0.18 | 1 | 136.07 | 1.30 | 0.26 |

***Table S9.*** *Weight day six minimal adequate model. Table consists of all factors tested in the linear mixed model with the weight from six-day-old testosterone treated and control nestlings as the dependent variable. Brood of origin (var = 1.04) and brood of rearing (0.00) were included as random factors. The estimate, degrees of freedom (numerator df and denominator df), the test statistic (F-value) and the significance (p-value) are given.*

| Weight day six | Estimate ± SE | Num. df | Denom. df | F-value | P-value |
|---|---|---|---|---|---|
| **Minimal adequate model** | | | | | |
| Intercept | 9.43 ± 0.30 | 1 | 41.56 | - | - |
| Treatment (T) | 0.14 ± 0.39 | 1 | 33.19 | 0.13 | 0.72 |
| Sex (male) | 0.41 ± 0.18 | 1 | 137.06 | 5.43 | **0.02** |
| **Dropped terms** | | | | | |
| Treatment (T) × Sex (male) | 0.22 ± 0.36 | 1 | 137.10 | 0.37 | 0.54 |

***Table S10.*** *Weight day eight minimal adequate model. Table consists of all factors tested in the linear mixed model with the weight from eight-day-old testosterone treated and control nestlings as the dependent variable. Brood of origin (var = 0.69) and brood of rearing (0.11) were included as random factors. The estimate, degrees of freedom (numerator df and denominator df), the test statistic (F-value) and the significance (p-value) are given.*

| Weight day eight | Estimate ± SE | Num. df | Denom. df | F-value | P-value |
|---|---|---|---|---|---|
| **Minimal adequate model** | | | | | |
| Intercept | 12.39 ± 0.29 | 1 | 49.62 | - | - |
| Treatment (T) | 0.18 ± 0.36 | 1 | 27.43 | 0.26 | 0.61 |
| Sex (male) | 0.06 ± 0.22 | 1 | 128.92 | 0.07 | 0.80 |
| **Dropped terms** | | | | | |
| Treatment (T) × Sex (male) | 0.54 ± 0.44 | 1 | 124.20 | 1.51 | 0.22 |

***Table S11.*** *Probability of begging day eight minimal adequate model. Table consists of all factors tested in the generalized linear mixed model with whether eight-day-old testosterone treated and control nestlings begged (yes/no) during a parental visit as the dependent variable. Nestling ID (var = 0.33), brood of origin (0.00) and brood of rearing (0.09) were included as random factors. The estimate, degrees of freedom (numerator df), the test statistic ($\chi^2$ value) and the significance (p-value) are given.*

| Probability of begging day eight | Estimate ± SE | Num. df | $\chi^2$ value | P-value |
|---|---|---|---|---|
| **Minimal adequate model** | | | | |
| Intercept | 0.46 ± 0.16 | 1 | - | - |
| Treatment (T) | -0.30 ± 0.15 | 1 | 3.99 | **< 0.05** |
| Sex (male) | -0.31 ± 0.15 | 1 | - | - |
| **Dropped terms** | | | | |
| Treatment (T) × Sex (male) | 0.29 ± 0.31 | 1 | 0.90 | 0.34 |

***Table S12.*** *Probability of getting fed day eight minimal adequate model. Table consists of all factors tested in the generalized linear mixed model with whether eight-day-old testosterone treated and control nestlings got fed (yes/no) during a parental visit as the dependent variable. Nestling ID (var = $5.45 \times 10^{-10}$), brood of origin (0.03) and brood of rearing (0.05) were included as random factors. The estimate, degrees of freedom (numerator df), the test statistic ($\chi^2$ value) and the significance (p-value) are given.*

| Probability of getting fed day eight | Estimate ± SE | Num. df | $\chi^2$ value | P-value |
|---|---|---|---|---|
| **Minimal adequate model** | | | | |
| Intercept | -1.39 ± 0.12 | 1 | - | - |
| Treatment (T) | -0.13 ± 0.12 | 1 | 1.02 | 0.31 |
| Sex (male) | -0.16 ± 0.10 | 1 | - | - |
| **Dropped terms** | | | | |
| Treatment (T) × Sex (male) | 0.12 ± 0.19 | 1 | 0.37 | 0.54 |

***Table S13.*** *Weight day 14 minimal adequate model. Table consists of all factors tested in the linear mixed model with the weight from 14-day-old testosterone treated and control nestlings as the dependent variable. Brood of origin (var = 0.56) and brood of rearing (0.19) were included as random factors. The estimate, degrees of freedom (numerator df and denominator df), the test statistic (F-value) and the significance (p-value) are given.*

| Weight day 14 | Estimate ± SE | Num. df | Denom. df | F-value | P-value |
|---|---|---|---|---|---|
| **Minimal adequate model** | | | | | |
| Intercept | 17.30 ± 0.22 | 1 | 55.38 | - | - |
| Treatment (T) | -0.48 ± 0.27 | 1 | 20.17 | 0.66 | 0.43 |
| Sex (male) | 0.41 ± 0.18 | 1 | 117.12 | 29.53 | < 0.001 |
| Treatment (T) × Sex (male) | 0.56 ± 0.25 | 1 | 110.24 | 5.24 | **0.02** |
| **Dropped terms** | | | | | |
| NA | - | - | - | - | - |

***Table S14.*** *Tarsus length day 14 minimal adequate model. Table consists of all factors tested in the linear mixed model with the tarsus length from 14-day-old testosterone treated and control nestlings as the dependent variable. Brood of origin (var = 0.05) and brood of rearing (0.01) were included as random factors. The estimate, degrees of freedom (numerator df and denominator df), the test statistic (F-value) and the significance (p-value) are given.*

| **Tarsus length day 14** | **Estimate ± SE** | **Num. df** | **Denom. df** | **F-value** | **P-value** |
|---|---|---|---|---|---|
| **Minimal adequate model** | | | | | |
| Intercept | 18.81 ± 0.08 | 1 | 48.48 | - | - |
| Treatment (T) | 0.18 ± 0.10 | 1 | 27.54 | 3.08 | 0.09 |
| Sex (male) | 0.55 ± 0.07 | 1 | 127.72 | 61.30 | **< 0.001** |
| **Dropped terms** | | | | | |
| Treatment (T) × Sex (male) | 0.09 ± 0.14 | 1 | 120.93 | 0.40 | 0.53 |

***Table S15.*** *Handling stress day 14 minimal adequate model. Table consists of all factors tested in the linear mixed model with the handling stress from 14-day-old testosterone treated and control nestlings as the dependent variable. Brood of origin (var = 1.06) and brood of rearing (0.06) were included as random factors. The estimate, degrees of freedom (numerator df and denominator df), the test statistic (F-value) and the significance (p-value) are given.*

| **Handling stress day 14** | **Estimate ± SE** | **Num. df** | **Denom. df** | **F-value** | **P-value** |
|---|---|---|---|---|---|
| **Minimal adequate model** | | | | | |
| Intercept | 1.07 ± 0.35 | 1 | 53.50 | - | - |
| Treatment (T) | 0.03 ± 0.44 | 1 | 32.85 | 0.006 | 0.94 |
| Sex (male) | -0.86 ± 0.27 | 1 | 124.63 | 10.38 | **0.002** |
| **Dropped terms** | | | | | |
| Treatment (T) × Sex (male) | 0.72 ± 0.52 | 1 | 122.21 | 1.90 | 0.17 |

**Appendix S6: Gene annotation and GO analysis**

***Table S16.*** *Number and annotation of CpG sites for which an effect of treatment was found (either in interaction with sex or corrected for sex).*

| | Significant CpGs | |
|---|---|---|
| | **Treatment × sex** | **Treatment + sex** |
| **Total** | 763 | 0 |
| **No annotation** | 113 | NA |
| **Annotated** | 650 | NA |
| **Promoter** | 138 | NA |
| **Of which TSS** | 18 | NA |
| **Gene body (exon or intron)** | 392 | NA |
| **Upstream** | 41 | NA |
| **Downstream** | 79 | NA |

***Table S17.*** *Sex-specific DMS in genes related to nervous system development and behaviour, focusing on sex-specific DMS within regulatory regions (promoter and TSS regions) of genes and sex-specific DMS that occurred in the same gene.*

| Gene | Findings (incl. relevant papers) |
|---|---|
| *FERMT2* | amyloid deposition (Chapuis et al., 2017), Alzheimer's disease, sex-specific gene expression (Fan et al., 2020) |
| *GJC2* | myelin maintenance, psychiatric history (Miguel-Hidalgo et al., 2017), sex-specific gene expression (Cao et al., 2013) |
| *GSX2* | oligodendrogenesis (Chapman et al., 2013) |
| *HOXD3* | posttraumatic stress disorder (Logue et al., 2015), sex-specific gene expression (Strawn et al., 2021) |
| *PTPRC* | Parkinson's Disease (Henderson et al., 2021) |
| *NEDD4L* | depression (Xu et al., 2020) |

***Table S18.*** *Sex-specific DMS in genes related to growth and metabolism, focusing on sex-specific DMS within regulatory regions (promoter and TSS regions) of genes and sex-specific DMS that occurred in the same gene.*

| Gene | Findings (incl. relevant papers) |
|---|---|
| *SEC22C,* | protein trafficking (Yamamoto et al., 2017) |
| *SLC12A2* | weight gain (Serão et al., 2013; Seabury et al., 2017) |
| *AMN* | embryonic development (Kalantry et al., 2001), nutrient transfer from the yolk sac to the developing chicken embryo (Bauer et al., 2013, 2020) |

***Table S19.*** *Sex-specific DMS in genes related to male fertility, focusing on sex-specific DMS within regulatory regions (promoter and TSS regions) of genes and sex-specific DMS that occurred in the same gene.*

| Gene | Findings (incl. relevant papers) |
|---|---|
| *ACYP2* | sperm motility (Gentiluomo et al., 2021), sex-specific gene expression (Wilhelm et al., 2016) |
| Intriguingly, we found fourteen sex-specific DMS in or near genes that are part of the solute carrier (SLC) group: *SLC4A1*, *SLC4A4*, *SLC6A5*, *SLC7A14*, *SLC12A2*, *SLC16A1*, *SLC20A2*, *SLC24A5*, *SLC30A2*, *SLC34A2*, *SLC39A11*, *SLC44A2* and *SLC45A3*. Literature is scarce on these specific genes. | |
| *SLC12A2* | spermatogenesis (Pace et al., 2000), castration (Xing et al., 2017), negative relationship between promoter methylation and expression of this gene has been found in rats (Lee et al., 2010), suggestion a causal relationship |
| *SLC16A1* | gene expression affected by experimentally increased yolk testosterone in male zebra finches (Bentz et al., 2021) |
| *SLC30A2* | SLC30 family might be important for testosterone synthesis and male reproduction (Chu et al., 2016) |

***Table S20.*** *Sex-specific DMS in genes related to oxidative stress and DNA repair, focusing on sex-specific DMS within regulatory regions (promoter and TSS regions) of genes and sex-specific DMS that occurred in the same gene.*

| Gene | Findings (incl. relevant papers) |
|---|---|
| *ACYP2* | stress-induced cell apoptosis (Kim et al., 2007), telomere length (Codd et al., 2013), sex-specific gene expression (Wilhelm et al., 2016) |
| *APEH* | oxidative stress (Tyler et al., 2021), sex-specific gene activity (Tyler et al., 2021) |
| *SWI5* | DNA recombination and repair (Akamatsu and Jasin, 2010; Kokabu et al., 2011; Yuan and Chen, 2011) |

**References Tables S17-S20**

Akamatsu, Y., and Jasin, M. (2010). Role for the Mammalian Swi5-Sfr1 Complex in DNA Strand Break Repair through Homologous Recombination. *PLOS Genetics* 6, e1001160. doi: 10.1371/journal.pgen.1001160

Bauer, R., Plieschnig, J. A., Finkes, T., Riegler, B., Hermann, M., and Schneider, W. J. (2013). The Developing Chicken Yolk Sac Acquires Nutrient Transport Competence by an Orchestrated Differentiation Process of Its Endodermal Epithelial Cells*. *Journal of Biological Chemistry* 288, 1088–1098. doi: 10.1074/jbc.M112.393090

Bauer, R., Tondl, P., and Schneider, W. J. (2020). A differentiation program induced by bone morphogenetic proteins 4 and 7 in endodermal epithelial cells provides the molecular basis for efficient nutrient transport by the chicken yolk sac. *Developmental Dynamics* 249, 222–236. doi: 10.1002/dvdy.129

Bentz, A. B., Niederhuth, C. E., Carruth, L. L., and Navara, K. J. (2021). Prenatal testosterone triggers long-term behavioral changes in male zebra finches: unravelling the neurogenomic mechanisms. *BMC Genomics* 22, 158. doi: 10.1186/s12864-021-07466-9

Cao, J., Wang, J., Dwyer, J. B., Gautier, N. M., Wang, S., Leslie, F. M., et al. (2013). Gestational nicotine exposure modifies myelin gene expression in the brains of adolescent rats with sex differences. *Transl Psychiatry* 3, e247–e247. doi: 10.1038/tp.2013.21

Chapman, H., Waclaw, R. R., Pei, Z., Nakafuku, M., and Campbell, K. (2013). The homeobox gene *Gsx2* controls the timing of oligodendroglial fate specification in mouse lateral ganglionic eminence progenitors. *Development* 140, 2289–2298. doi: 10.1242/dev.091090

Chapuis, J., Flaig, A., Grenier-Boley, B., Eysert, F., Pottiez, V., Deloison, G., et al. (2017). Genome-wide, high-content siRNA screening identifies the Alzheimer's genetic risk factor FERMT2 as a major modulator of APP metabolism. *Acta Neuropathol* 133, 955–966. doi: 10.1007/s00401-016-1652-z

Chu, Q., Chi, Z.-H., Zhang, X., Liang, D., Wang, X., Zhao, Y., et al. (2016). A potential role for zinc transporter 7 in testosterone synthesis in mouse Leydig tumor cells. *International Journal of Molecular Medicine* 37, 1619–1626. doi: 10.3892/ijmm.2016.2576

Codd, V., Nelson, C. P., Albrecht, E., Mangino, M., Deelen, J., Buxton, J. L., et al. (2013). Identification of seven loci affecting mean telomere length and their association with disease. *Nat Genet* 45, 422-427e2. doi: 10.1038/ng.2528

Fan, C. C., Banks, S. J., Thompson, W. K., Chen, C.-H., McEvoy, L. K., Tan, C. H., et al. (2020). Sex-dependent autosomal effects on clinical progression of Alzheimer's disease. *Brain* 143, 2272–2280. doi: 10.1093/brain/awaa164

Gentiluomo, M., Luddi, A., Cingolani, A., Fornili, M., Governini, L., Lucenteforte, E., et al. (2021). Telomere Length and Male Fertility. *International Journal of Molecular Sciences* 22, 3959. doi: 10.3390/ijms22083959

Henderson, A. R., Wang, Q., Meechoovet, B., Siniard, A. L., Naymik, M., De Both, M., et al. (2021). DNA Methylation and Expression Profiles of Whole Blood in Parkinson's Disease. *Frontiers in Genetics* 12. Available at: https://www.frontiersin.org/article/10.3389/fgene.2021.640266 (Accessed May 19, 2022).

Kalantry, S., Manning, S., Haub, O., Tomihara-Newberger, C., Lee, H.-G., Fangman, J., et al. (2001). The amnionless gene, essential for mouse gastrulation, encodes a visceral-endoderm–specific protein with an extracellular cysteine-rich domain. *Nat Genet* 27, 412–416. doi: 10.1038/86912

Kim, J. W., Kwon, O. Y., and Kim, M. H. (2007). Differentially expressed genes and morphological changes during lengthened immobilization in rat soleus muscle. *Differentiation* 75, 147–157. doi: 10.1111/j.1432-0436.2006.00118.x

Kokabu, Y., Murayama, Y., Kuwabara, N., Oroguchi, T., Hashimoto, H., Tsutsui, Y., et al. (2011). Fission Yeast Swi5-Sfr1 Protein Complex, an Activator of Rad51 Recombinase, Forms an Extremely Elongated Dogleg-shaped Structure *. *Journal of Biological Chemistry* 286, 43569–43576. doi: 10.1074/jbc.M111.303339

Lee, H.-A., Hong, S.-H., Kim, J.-W., and Jang, I.-S. (2010). Possible involvement of DNA methylation in NKCC1 gene expression during postnatal development and in response to ischemia. *Journal of Neurochemistry* 114, 520–529. doi: 10.1111/j.1471-4159.2010.06772.x

Logue, M. W., Smith, A. K., Baldwin, C., Wolf, E. J., Guffanti, G., Ratanatharathorn, A., et al. (2015). An analysis of gene expression in PTSD implicates genes involved in the glucocorticoid receptor pathway and neural responses to stress. *Psychoneuroendocrinology* 57, 1–13. doi: 10.1016/j.psyneuen.2015.03.016

Miguel-Hidalgo, J. J., Hall, K. O., Bonner, H., Roller, A. M., Syed, M., Park, C. J., et al. (2017). MicroRNA-21: Expression in oligodendrocytes and correlation with low myelin mRNAs in depression and alcoholism. *Progress in Neuro-Psychopharmacology and Biological Psychiatry* 79, 503–514. doi: 10.1016/j.pnpbp.2017.08.009

Pace, A. J., Lee, E., Athirakul, K., Coffman, T. M., O'Brien, D. A., and Koller, B. H. (2000). Failure of spermatogenesis in mouse lines deficient in the $Na^{+}$-$K^{+}$-$2Cl^{-}$ cotransporter. *J Clin Invest* 105, 441–450. doi: 10.1172/JCI8553

Seabury, C. M., Oldeschulte, D. L., Saatchi, M., Beever, J. E., Decker, J. E., Halley, Y. A., et al. (2017). Genome-wide association study for feed efficiency and growth traits in U.S. beef cattle. *BMC Genomics* 18, 386. doi: 10.1186/s12864-017-3754-y

Serão, N. V., González-Peña, D., Beever, J. E., Faulkner, D. B., Southey, B. R., and Rodriguez-Zas, S. L. (2013). Single nucleotide polymorphisms and haplotypes associated with feed efficiency in beef cattle. *BMC Genet* 14, 94. doi: 10.1186/1471-2156-14-94

Strawn, M., Moraes, J. G. N., Safranski, T. J., and Behura, S. K. (2021). Sexually Dimorphic Transcriptomic Changes of Developing Fetal Brain Reveal Signaling Pathways and Marker Genes of Brain Cells in Domestic Pigs. *Cells* 10, 2439. doi: 10.3390/cells10092439

Tyler, K., Geilman, S., Bell, D. M., Taylor, N., Honeycutt, S. C., Garrett, P. I., et al. (2021). Acyl Peptide Enzyme Hydrolase (APEH) activity is inhibited by lipid metabolites and peroxidation products. *Chemico-Biological Interactions* 348, 109639. doi: 10.1016/j.cbi.2021.109639

Wilhelm, C. J., Hashimoto, J. G., Roberts, M. L., Bloom, S. H., Andrew, M. R., and Wiren, K. M. (2016). Astrocyte Dysfunction Induced by Alcohol in Females but Not Males. *Brain Pathology* 26, 433–451. doi: 10.1111/bpa.12276

Xing, B., Bai, X., Guo, H., Chen, J., Hua, L., Zhang, J., et al. (2017). Long non-coding RNA analysis of muscular responses to testosterone deficiency in Huainan male pigs. *Animal Science Journal* 88, 1451–1456. doi: 10.1111/asj.12777

Xu, J., Guo, C., Liu, Y., Wu, G., Ke, D., Wang, Q., et al. (2020). Nedd4l downregulation of NRG1 in the mPFC induces depression-like behaviour in CSDS mice. *Transl Psychiatry* 10, 249. doi: 10.1038/s41398-020-00935-x

Yamamoto, Y., Yurugi, C., and Sakisaka, T. (2017). The number of the C-terminal transmembrane domains has the potency to specify subcellular localization of Sec22c. *Biochemical and Biophysical Research Communications* 487, 388–395. doi: 10.1016/j.bbrc.2017.04.071

Yuan, J., and Chen, J. (2011). The Role of the Human SWI5-MEI5 Complex in Homologous Recombination Repair *. *Journal of Biological Chemistry* 286, 9888–9893. doi: 10.1074/jbc.M110.207290

***Table S21.*** *Enriched GO terms for the ontology biological process with significant FDR q-values for the genes associated with antagonistic DMS. The number of genes in the target list annotated to the particular GO category and the number of genes in the background list annotated to a certain GO category are shown in the x and n columns, respectively. The total number of (recognised) genes in the target list was 198 and the total number of genes in the background list was 9303.*

| GO term | Description | FDR q-value | x | n |
|---|---|---|---|---|
| GO:0050796 | regulation of insulin secretion | 0.02 | 8 | 111 |
| GO:0030072 | peptide hormone secretion | 0.02 | 9 | 163 |
| GO:0045685 | regulation of glial cell differentiation | 0.02 | 5 | 58 |
| GO:0071692 | protein localization to extracellular region | 0.02 | 12 | 237 |
| GO:0002790 | peptide secretion | 0.02 | 9 | 165 |
| GO:0090087 | regulation of peptide transport | 0.02 | 8 | 133 |
| GO:0002791 | regulation of peptide secretion | 0.02 | 8 | 132 |
| GO:0008360 | regulation of cell shape | 0.02 | 7 | 105 |
| GO:0007043 | cell-cell junction assembly | 0.02 | 7 | 105 |
| GO:0035592 | establishment of protein localization to extracellular region | 0.02 | 12 | 234 |
| GO:0009306 | protein secretion | 0.02 | 12 | 234 |
| GO:0046883 | regulation of hormone secretion | 0.02 | 9 | 173 |
| GO:0090276 | regulation of peptide hormone secretion | 0.02 | 8 | 131 |
| GO:0034284 | response to monosaccharide | 0.02 | 9 | 136 |
| GO:0045216 | cell-cell junction organization | 0.02 | 8 | 143 |
| GO:0050708 | regulation of protein secretion | 0.02 | 9 | 170 |
| GO:0099175 | regulation of postsynapse organization | 0.02 | 5 | 66 |
| GO:0021510 | spinal cord development | 0.02 | 6 | 88 |
| GO:0007156 | homophilic cell adhesion via plasma membrane adhesion molecules | 0.02 | 6 | 79 |
| GO:0002793 | positive regulation of peptide secretion | 0.02 | 5 | 67 |
| GO:0090277 | positive regulation of peptide hormone secretion | 0.02 | 5 | 67 |
| GO:0009743 | response to carbohydrate | 0.02 | 9 | 147 |
| GO:0015833 | peptide transport | 0.03 | 9 | 180 |
| GO:0014013 | regulation of gliogenesis | 0.03 | 5 | 71 |
| GO:0007160 | cell-matrix adhesion | 0.03 | 8 | 154 |
| GO:0048483 | autonomic nervous system development | 0.03 | 5 | 44 |
| GO:0050773 | regulation of dendrite development | 0.03 | 5 | 72 |
| GO:0032024 | positive regulation of insulin secretion | 0.03 | 5 | 50 |
| GO:0048813 | dendrite morphogenesis | 0.03 | 6 | 104 |
| GO:0030183 | B cell differentiation | 0.03 | 5 | 75 |
| GO:0090101 | negative regulation of transmembrane receptor protein serine/threonine kinase signaling pathway | 0.03 | 6 | 105 |

***Table S21 (continued).***

| GO term | Description | FDR q-value | x | n |
|---|---|---|---|---|
| GO:0048706 | embryonic skeletal system development | 0.03 | 6 | 103 |
| GO:0097696 | receptor signaling pathway via STAT | 0.03 | 5 | 78 |
| GO:0030073 | insulin secretion | 0.03 | 9 | 133 |
| GO:0035270 | endocrine system development | 0.03 | 6 | 108 |
| GO:0007605 | sensory perception of sound | 0.04 | 6 | 111 |
| GO:0045598 | regulation of fat cell differentiation | 0.04 | 5 | 84 |
| GO:0050714 | positive regulation of protein secretion | 0.04 | 5 | 87 |
| GO:0001837 | epithelial to mesenchymal transition | 0.05 | 6 | 119 |
| GO:0009746 | response to hexose | 0.05 | 9 | 131 |
| GO:0046887 | positive regulation of hormone secretion | 0.05 | 5 | 90 |

***Table S22.*** *Enriched GO terms for the ontologies cellular component and molecular function with significant FDR q-values for the genes associated with antagonistic DMS. The number of genes in the target list annotated to the particular GO category and the number of genes in the background list annotated to a certain GO category are shown in the x and n columns, respectively. The total number of (recognised) genes in the target list was 198 and the total number of genes in the background list was 9303.*

| GO term | Description | FDR q-value | x | n |
|---|---|---|---|---|
| **Cellular component** | | | | |
| GO:0005903 | brush border | 0.02 | 5 | 66 |
| GO:0044306 | neuron projection terminus | 0.03 | 6 | 95 |
| GO:0019898 | extrinsic component of membrane | 0.03 | 10 | 187 |
| GO:0005912 | adherens junction | 0.03 | 7 | 133 |
| GO:0043679 | axon terminus | 0.04 | 5 | 84 |
| GO:0016363 | nuclear matrix | 0.04 | 5 | 85 |
| **Molecular function** | | | | |
| GO:0016849 | phosphorus-oxygen lyase activity | 0.02 | 5 | 51 |
| GO:0001221 | transcription coregulator binding | 0.02 | 6 | 90 |

***Table S23.*** *Enriched GO terms for the ontologies biological process, cellular component and molecular function with significant FDR q-values for the genes associated with female-specific and male-specific DMS. The number of genes in the target list annotated to the particular GO category and the number of genes in the background list annotated to a certain GO category are shown in the x and n columns, respectively. The total number of (recognised) genes in the female-specific target list was 120, the total number in the male-specific target list was 126 and the total number in the background list was 9303.*

| **DMS** | **GO term** | **Description** | **FDR q-value** | **x** | **n** |
|---|---|---|---|---|---|
| **Biological process** | | | | | |
| Female-specific | GO:0021510 | spinal cord development | 0.001 | 7 | 88 |
| Female-specific | GO:0001764 | neuron migration | 0.001 | 8 | 133 |
| Female-specific | GO:0021766 | hippocampus development | 0.004 | 5 | 66 |
| Female-specific | GO:0061351 | neural precursor cell proliferation | 0.004 | 6 | 111 |
| Female-specific | GO:0048709 | oligodendrocyte differentiation | 0.004 | 5 | 76 |
| Female-specific | GO:0021761 | limbic system development | 0.007 | 5 | 92 |
| Female-specific | GO:0030048 | actin filament-based movement | 0.007 | 5 | 90 |
| **Cellular component** | | | | | |
| Female-specific | GO:0016363 | nuclear matrix | 0.002 | 6 | 85 |
| Female-specific | GO:0034399 | nuclear periphery | 0.003 | 6 | 98 |
| **Molecular function** | | | | | |
| Female-specific | - | - | - | - | - |
| **Biological process** | | | | | |
| Male-specific | GO:0048704 | embryonic skeletal system morphogenesis | 0.002 | 6 | 74 |
| Male-specific | GO:0051216 | cartilage development | 0.003 | 8 | 150 |
| Male-specific | GO:0048706 | embryonic skeletal system development | 0.004 | 7 | 103 |
| Male-specific | GO:0006109 | regulation of carbohydrate metabolic process | 0.007 | 6 | 115 |
| Male-specific | GO:0031214 | biomineral tissue development | 0.008 | 6 | 111 |
| Male-specific | GO:0035113 | embryonic appendage morphogenesis | 0.01 | 5 | 98 |
| Male-specific | GO:0030326 | embryonic limb morphogenesis | 0.01 | 5 | 98 |
| Male-specific | GO:0001708 | cell fate specification | 0.01 | 5 | 97 |
| **Cellular component** | | | | | |
| Male-specific | - | - | - | - | - |
| **Molecular function** | | | | | |
| Male-specific | GO:0001221 | transcription coregulator binding | 0.01 | 5 | 90 |
